# Single-cell-resolved interspecies comparison identifies a shared inflammatory axis and a dominant neutrophil-endothelial program in severe COVID-19

**DOI:** 10.1101/2023.08.25.551434

**Authors:** Stefan Peidli, Geraldine Nouailles, Emanuel Wyler, Julia M. Adler, Sandra Kunder, Anne Voß, Julia Kazmierski, Fabian Pott, Peter Pennitz, Dylan Postmus, Luiz Gustavo Teixeira Alves, Christine Goffinet, Achim D. Gruber, Nils Blüthgen, Martin Witzenrath, Jakob Trimpert, Markus Landthaler, Samantha D. Praktiknjo

## Abstract

Key issues for research of COVID-19 pathogenesis are the lack of biopsies from patients and of samples at the onset of infection. To overcome these hurdles, hamsters were shown to be useful models for studying this disease. Here, we further leveraged the model to molecularly survey the disease progression from time-resolved single-cell RNA-sequencing data collected from healthy and SARS-CoV-2-infected Syrian and Roborovski hamster lungs. We compared our data to human COVID-19 studies, including BALF, nasal swab, and post-mortem lung tissue, and identified a shared axis of inflammation dominated by macrophages, neutrophils, and endothelial cells, which we show to be transient in Syrian and terminal in Roborovski hamsters. Our data suggest that, following SARS-CoV-2 infection, commitment to a type 1 or type 3-biased immunity determines moderate versus severe COVID-19 outcomes, respectively.

**One-Sentence Summary:** Activation of different immunological programs upon SARS-CoV-2 infection determines COVID-19 severity.

## Introduction

For coronavirus disease 2019 (COVID-19), caused by infection with severe acute respiratory syndrome coronavirus 2 (SARS-CoV-2), a range of cellular and molecular processes have been associated with the first acute disease phase (*1, 2*). Among them, particularly innate immunity responses have been shown to be essential in modulating severe / critical COVID-19, ultimately leading to an overly exacerbated inflammatory state and the production of pro-inflammatory cytokines (*3-9*). This has been extensively described for pulmonary macrophages (*10, 11*) but also for neutrophils, which contribute to disease severity through thrombotic complications caused by NETosis (*12, 13*).

Animal models have been essential in the study of COVID-19 (*14, 15*). Hamsters, which, in contrast to mice, can be readily infected by the same SARS-CoV-2 variants as humans, have been particularly useful for the molecular investigation of COVID-19-related pathomechanisms (*16, 17*). A key advantage of animal models is that reactions to the infection can be studied from the earliest time point after infection. This is in contrast to patient samples, for which sampling usually happens the earliest about a week after infection due to incubation times and study enrollment. Furthermore, animal models allow for investigation of samples which are not accessible in humans due to medical and ethical constraints, such as whole lung tissue, thereby making it possible to identify early processes of innate immunity that are fundamental in driving the initial onset and set the course towards mild or severe COVID-19.

Here, we provide a comprehensive survey of the dynamic cellular and molecular pulmonary landscape underlying different COVID-19 outcomes. Specifically, based on the observations that Syrian hamsters (*Mesocricetus auratus*) experience a moderate course of disease, whereas Roborovski hamsters (*Phodopus roborovskii*) suffer from severe to lethal COVID-19 (*16, 18*), we established a time-resolved resource of single-cell RNA-sequencing (scRNA-seq) data collected from healthy and SARS-CoV-2-infected lung tissue of both hamster species that we extensively compared to published human data. Using a range of analysis methods, including post-hoc interpretation methods in cell type-specific diffusion map latent spaces, we found neutrophils and endothelial cells to be instrumental in regulating severe courses of COVID-19, and identify expression programs that potentially orchestrate progressive endothelial damage.

## Results

### Tracking cellular changes throughout SARS-CoV-2 infection

To identify the cells and early molecular markers that regulate COVID-19 disease severity, we jointly analyzed pulmonary single-cell RNA (scRNA) time-course data from two different hamster species modeling moderate and severe COVID-19. Specifically, we reprocessed our previously published data for Syrian hamsters (*19*) and generated de novo Roborovski hamster scRNA-sequencing (seq) datasets for this study which we further compared to publicly-available human data (*20-22*) (Fig. 1A).

**Fig. 1.**
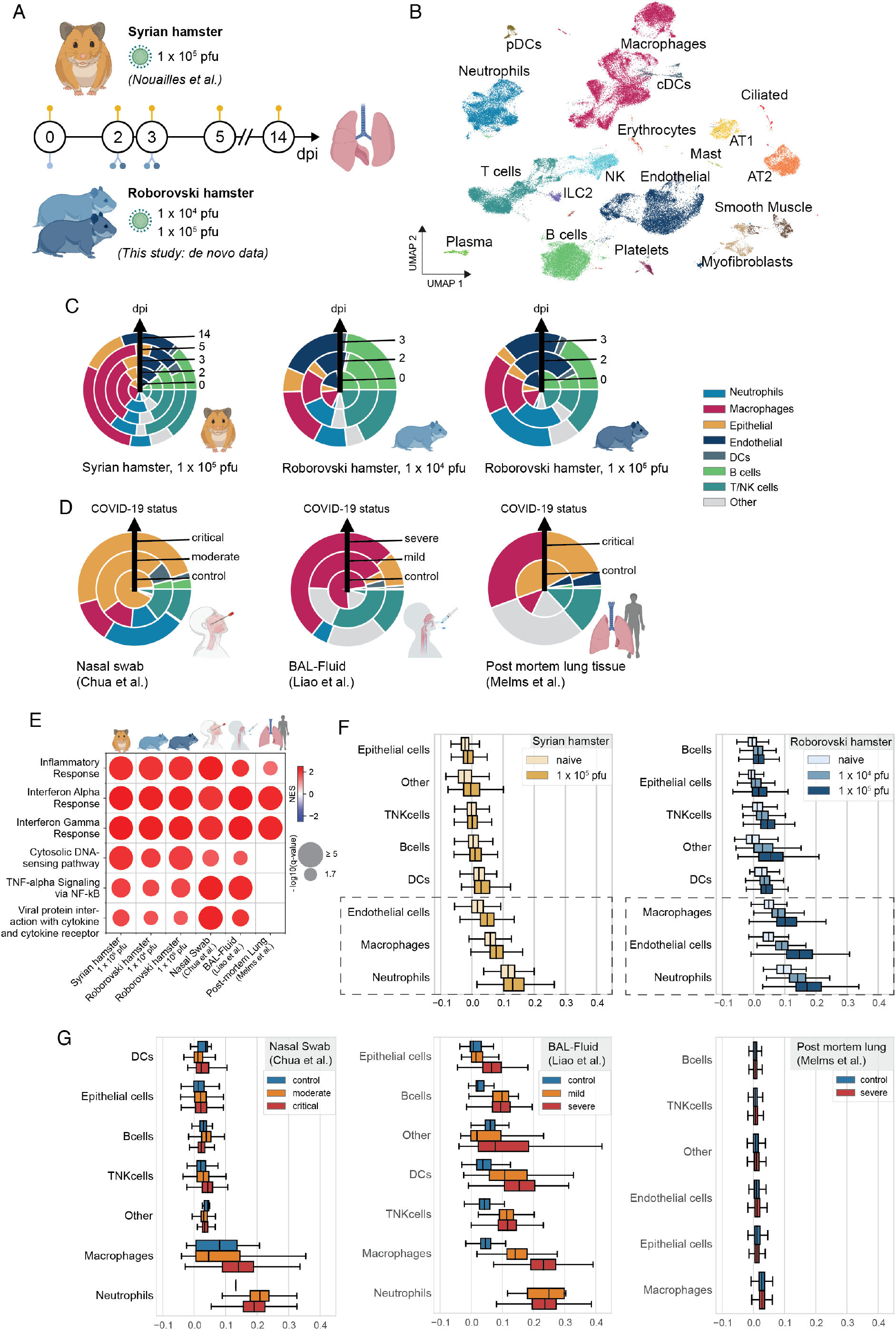
The single-cell landscape of SARS-CoV-2 infected lungs. (**A**) Study design outlining hamster data used in this study. Syrian and Roborovski hamsters were challenged with SARS-CoV-2 (1×10^4^ pfu or 1×10^5^ pfu SARS-CoV-2 Variant B.1, as indicated), and scRNA-seq data collected from lung tissue 0 (naïve / control), 2, and 3 days post infection (dpi) for Roborovski hamsters treated with two different infection doses, and additionally 5 and 14 dpi for Syrian hamsters. Syrian hamster dataset previously published in Nouailles et al. (*19*); n=3 biological replicates per sample point, 30 hamsters in total. (**B**) Integrated UMAP embedding of Syrian and Roborovski scRNA-seq samples with individual cells colored by cell type (**C**) Cell proportions per hamster species and virus dose colored by cell type. Each ring corresponds to a time point after infection in ascending order from the center outwards. (**D**) As in C, but for human datasets and with rings corresponding to COVID-19 status. (**E**) Selected significant gene sets from gene set enrichment analyses (GSEA) performed on genes differentially expressed in infected (2 and 3 dpi) vs. control samples across datasets. Color corresponds to normalized enrichment scores (NES) and dot size to -log10(q-value). Gene sets from KEGG (*54-56*) and MSigDB Hallmarks (*57, 58*). (**F**) Inflammatory response scores (Hallmark gene set, MSigDB; see Methods) across cell types in Syrian and Roborovski hamsters in infected (2 and 3 dpi) and naïve samples. (**G**) As in F, but for human datasets split by COVID-19 status. (**F, G**) Boxplots: box represents quartiles; line represents median; whiskers represent quartiles plus 1.5 times interquartile range added; outliers not shown.

Syrian hamsters develop moderate COVID-19 and recover from infection (*23*); therefore, analysis time points range from naïve animals (0 days post infection (dpi)), early onset (2 dpi) over acute phase of infection (3 dpi and 5 dpi) to recovery at 14 dpi. Roborovski hamsters develop an overall more severe course of disease, with fulminant pneumonia when infected with 1×10^5^ pfu and with less striking but still considerable COVID-19-like disease when infected with 1×10^4^ pfu of SARS-CoV-2. Time course data comprise naïve (0 dpi) and acute phase (2 and 3 dpi), as severe course hamsters do not recover from disease and reach humane end-point criteria from 3 dpi onwards. Objective clinical signs of SARS-CoV-2 infection are dominated by weight loss in Syrian hamsters, but include drop of body temperature and drastically reduced general condition in Roborovski hamsters that experience a severe course of infection (fig. S1A, (*18, 19*)). Single-cell hamster datasets displayed high quality and little variation across replicates (fig. S1B, C), enabling us to identify all major cell types of the lungs (Fig. 1B), including the primary target of SARS-CoV-2: Alveolar type 2 (AT2) cells (*24*). In addition, we established a customized data preprocessing pipeline (see Methods) that allows the recovery of many neutrophils that are typically discarded during filtering steps. Virus sequences were enriched in professional phagocytes (macrophages, neutrophils) (fig. S1D).

In order to relate the cellular composition of infected hamster lungs to human COVID-19 patients, we collected public scRNA-seq data from human nasal swab (*20*), bronchoalveolar lavage (BAL) fluid (*21*), and post-mortem lung tissue (*22*) samples. Direct comparison was hampered by different sample type, time of sampling, and variable quality of human single-cell datasets generally containing a lower complexity of cell types. Nevertheless, we observed shared COVID-19-specific trends. This particularly includes neutrophils, for which the proportion increased after infection across species and sample type with the exception of post-mortem lung tissue, for which no neutrophils could be detected (Fig. 1C, D). The increase in neutrophils was strongest in critical / severe human patients and Roborovski hamsters irrespective of virus dose, compared to a weaker rise in patients with mild / moderate COVID-19 and Syrian hamsters. Furthermore, T and NK cell proportions increased in human nasal swab and BAL-fluid samples after infection and Syrian hamsters towards 5 dpi. This was not observed in Roborovski hamsters, which might be due to the lack of corresponding samples from later stages of infection.

Beyond changes in cell frequency, we sought to identify cell types most strongly reacting to the infection. We therefore performed a global pathway enrichment analysis and found inflammatory genes to be highly enriched following SARS-CoV-2 infection throughout all hamster and human datasets (Fig. 1E, fig. S1E, see Methods). In both hamster species, neutrophils showed the strongest inflammatory activity among all cell types at the early onset and acute phase of infection (Fig. 1F), followed by macrophages. This also applied to human BAL-fluid and nasal swab (Fig. 1G). Moreover, we found that endothelial cells were strongly affected by the infection in lung tissues, with Roborovski hamsters having a markedly higher inflammation score than their Syrian hamster counterparts. Unlike BAL-fluid and nasal swab, human post-mortem lung tissue contained endothelial cells. However, data quality was overall low in this sample type with absence or extremely weak presence of detectable molecular signals. As a result, the feasibility of conducting further molecular investigation using this sample type was limited (Fig. 1G).

### SARS-CoV-2 infection causes cellular disbalance of innate immune cells

Innate immune cells in patients and animal models reacted strongly to SARS-CoV-2 infection, both in terms of composition and inflammatory gene expression profiles. We thus investigated these cells at higher resolution in lungs (Fig. 2A). By reintegrating and subclustering the respective cell populations, we were able to identify neutrophils and natural killer (NK) cells using canonical marker expression (*19*), in addition to four different subtypes of monocytes / macrophages including interstitial (IM), monocytic (MoM), alveolar macrophages (AM) and *Treml4*^+^ monocytes.

**Fig. 2.**
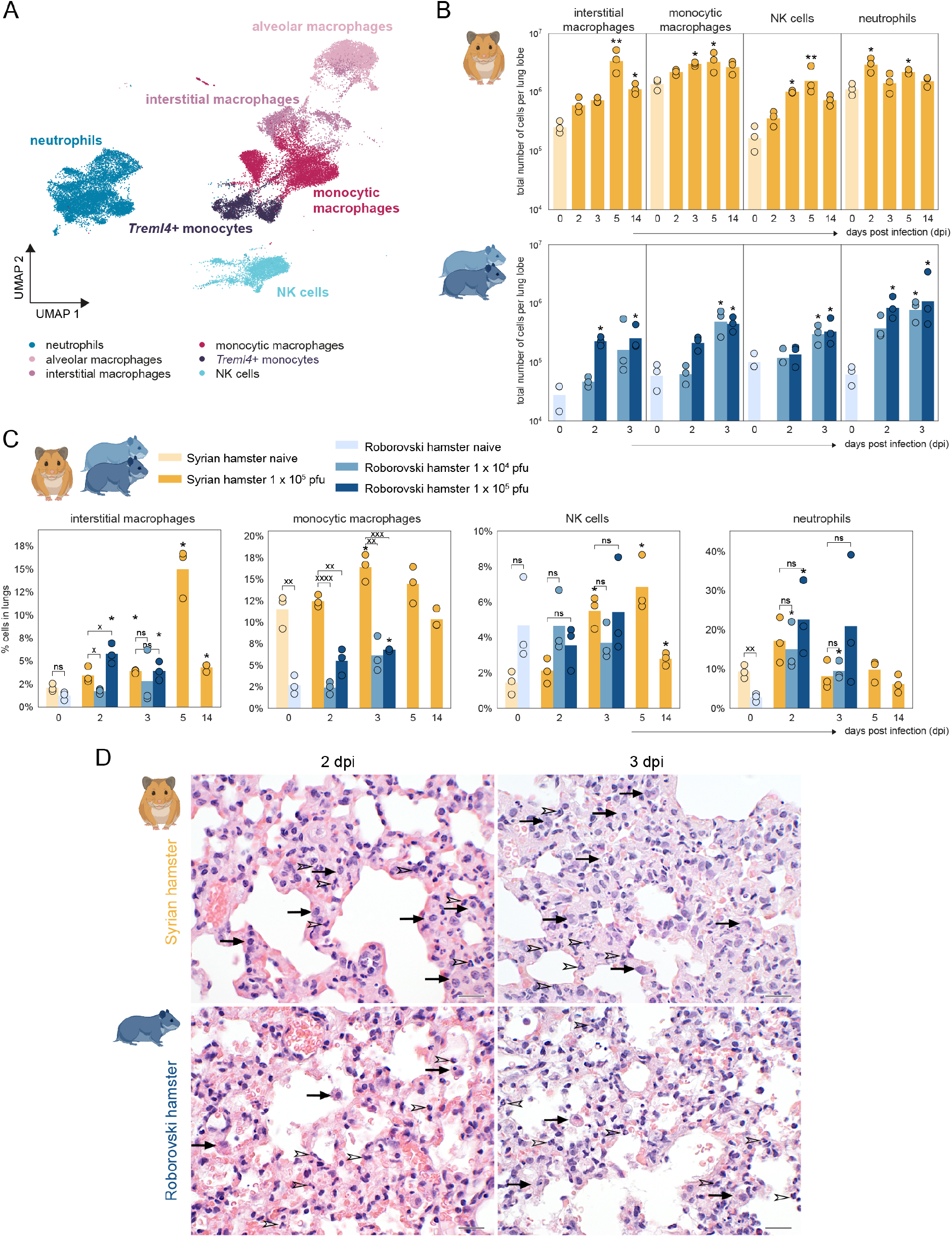
Disproportionate innate immune cell response throughout SARS-CoV-2 infection. (**A**) UMAP embedding of innate immune cells. Subclustering identified four distinct types of macrophages. (**B**) Total number of cells per selected cell type in lung lobe across days post infection (dpi), split by Syrian and Roborovski hamsters and additionally by virus dose for the latter, indicated by color. Test for difference in distribution across samples (n=3; one-way ANOVA with Dunn post-hoc test) against corresponding uninfected samples. *: < 0.05, ** < 0.01. (**C**) Relative numbers of cell types as shown in Fig. 2B. Additional test for difference in distribution (as described in (B)) across hamster types within time points. x < 0.05, xx < 0.01, xxx < 0.001, xxxx < 0.0001. (**D**) Histopathological comparison showing more cells compatible with macrophages (black arrows) and similar numbers of neutrophils (open arrowheads) in Syrian (top) than Roborovski (bottom) hamster lungs at 2 (left) and 3 (right) dpi, both infected with 1×10^5^ pfu SARS-CoV-2. H&E stain, original magnification = 600x, scale bars = 20 μm.

We forewent direct comparison of cell numbers per lobe between Syrian and Roborovski hamsters due to different animal and consequently lungs sizes (Fig. 2B). Overall, pulmonary numbers of IMs, MoMs, NK cells, and neutrophils increased significantly in both species during infection. In Syrian hamsters, IMs, MoMs and NK cell numbers peaked at 5 dpi, whereas the highest neutrophil numbers were detected at 2 dpi (Fig. 2B). Due to the short observation window, resolution of innate cellular responses was not observed in Roborovski hamsters (Fig. 2B). To further identify dominating immune cell types and for direct comparison between related species, we tracked cellular proportions which reflect local proliferation and cellular influx (Fig. 2C). IMs proportions increased fastest (2 dpi, ∼ 6%) upon high-dose infection of Roborovski hamsters, however, remained at similar levels by 3 dpi. In Syrian hamsters, IMs peaked by far highest with ∼15% amongst lung cells at 5 dpi, yet resolved close to baseline levels by 14 dpi. MoMs proportions were high (∼ 11%) in naïve (0 dpi) Syrian hamsters, yet significantly increased to ∼16% by 3 dpi. Similarly, MoMs increased significantly upon high-dose infection in Roborovski hamsters from ∼ 2% to ∼ 7% by 3 dpi. In contrast, the proportion pattern for NK cells and neutrophils differed between species. While NK cells significantly increased by 2-fold at 3 dpi compared to 0 dpi and peaked with ∼ 7% at 5 dpi in Syrian hamsters, SARS-CoV-2 infection did not trigger an increase in NK cell frequencies in Roborovski hamsters. Frequencies of neutrophils, however, increased significantly in Roborovski hamsters from ∼ 2% in naïve animals to ∼15% and ∼20% 2 days upon low and high-dose infection, respectively. Notably, only upon high-dose infection Roborovski hamsters maintained high neutrophil frequencies at 3 dpi. Syrian hamsters had higher baseline proportion of neutrophils (∼10%). However, neutrophils only modestly increased to ∼ 17% at 2 dpi, and started resolving already at 3 dpi to baseline levels (Fig. 2C). In contrast, AMs and *Treml4*^*+*^ monocytes did not react as strongly to infection (fig. S2A, B).

Histopathological examination revealed slightly higher numbers of cells consistent with macrophages in Syrian compared to Roborovski hamsters at 2 and 3 dpi despite similarly high infectious dose (Fig. 2D). Alveolar wall necrosis in Roborovski hamsters was generally more apparent when compared to Syrian hamsters at both time points, particularly at 3 dpi. Notably, Syrian hamsters developed first signs of epithelial cell proliferation and tissue regeneration at 3 dpi, resulting in a markedly higher density of parenchymal cells and thicker alveolar walls when compared to the more damaged and necrotic alveolar walls in Roborovski hamsters (Fig. 2D).

### Different cell-mediated immune programs are activated upon infection

The observed differences in innate immune cell composition upon infection indicate that different immunological programs are activated in Syrian and Roborovski hamsters. Innate and adaptive cell-mediated effector immunity can be discriminated into three types (*25*). Type 1 immunity is directed towards intracellular pathogens and achieves this primarily through the activation of macrophages and cytotoxic effector cells. Type 2 immunity targets helminths and stimulates mucus production and activation of mast cells, eosinophils, and basophils. Type 3 immunity is directed towards extracellular pathogens such as bacteria or fungi. Its effector modules trigger epithelial barrier defense and neutrophil recruitment.

In contrast to accessory cells, such as neutrophils and macrophages, which lack intrinsic specificity and are recruited to participate in a given immune response, NK cells, ILCs, and T cells express type-defining transcription factors. We therefore examined these cell types in the single-cell data in order to identify the dominant type of immunity (Fig. 3A, B). In accordance with preferential NK cell recruitment, we observed an increase of type 1 immunity-associated transcription factor (TF) expression (*Eomes, Tbx21*) (*26*) in lungs of Syrian hamsters, which was largely absent or weaker in Roborovski hamsters (Fig. 3B). The expression of these markers was confined to NK cells and partially to CD4^+^ T cells (Fig. 3A). Next, we visualized selected effector molecules, key cytokine and immune regulatory genes across time and all cell types to further evaluate the induced immune type (Fig. 3C, fig. S3A). The expression of *Eomes* clearly increased in Syrian until 5 dpi, while this trend was entirely absent in Roborovski hamsters. Additionally, the expression of typical cytotoxic effector molecules such as *Gzma, Prf1* and key type 1 cytokine *Ifng* were following similar trends in Syrian hamsters, while there was no consistent upregulation of *Gzma* and *Ifng* in lungs of Roborovski hamsters (Fig. 3C). Despite the predominant neutrophil response in Roborovski hamsters, the primary TF associated with type 3 immunity (*Rorc*) displayed no clear upregulation over time and all cells in high-dose infected Roborovski hamsters (Fig. 3C). However, focusing on the NK, T, and ILC clusters, *Rorc* expression significantly decreased at 5 dpi in Syrian hamsters and showed a trend towards upregulation at 2 dpi in high-dose and at 3 dpi in low-dose infected Roborovski hamsters (Fig. 3B). The type 2 immunity transcription factor GATA-3 was expressed in ILC2 and endothelial cell subsets as expected (*27, 28*) (Fig. 3A), while no upregulation upon infection was observed in the T/NK/ILC subset (Fig. 3B). Suppressor of cytokine signaling (SOCS) proteins regulate innate and adaptive immunity and can thereby shape initiation and maintenance of immune type. Broadly, SOCS1 suppresses type 1 and SOCS3 type 3 immunity (*29*). We found that at 2 dpi, *Socs1, Stat1* and *Il6* were strongly upregulated in Roborovski hamsters upon high-dose infection (Fig. 3C, fig. S3A). In Syrian hamsters, *Socs1* and *Socs3* were both upregulated at the peak cellular response at 5 dpi (Fig. 3C), and expression of *Il12a, Stat4*, and *Stat3* was generally higher compared to Roborovski hamsters (fig. S3A). Similarly, upon infection, *Ifng, Gzma*, and *Prf1* displayed uniform upregulation in Syrian hamsters in the T/NK/ILC subset, while *Stat1* and *Socs1* showed highest upregulation in high-dose infected Roborovski hamsters. *Stat4* and *Stat3*, however, displayed no clear trend towards upregulation or a specific hamster species bias in the T/NK/ILC subset (fig. S3B). In summary, the underlying gene expression profiles in Syrian and Roborovski hamsters broadly match the observed cellular effector responses, reinforcing a type 1 in Syrian versus type 3 bias in Roborovski hamsters.

**Fig. 3.**
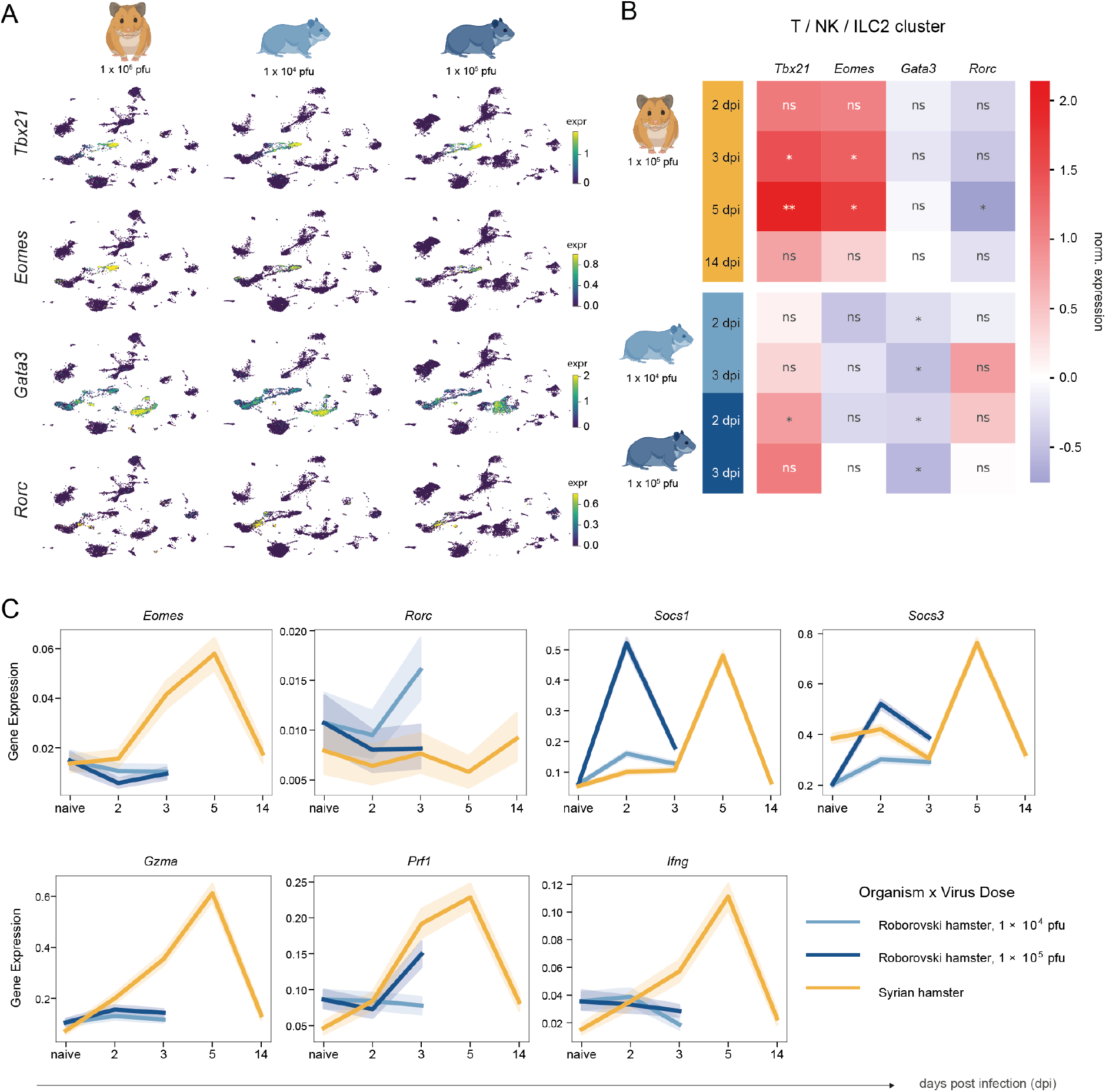
Roborovski and Syrian hamsters commit to different types of immunity. (**A**) Integrated UMAP embeddings as in Fig. 1B with normalized gene expressions of selected immunity-related transcription factors at 2 and 3 dpi for shown hamster species and virus dose. (**B**) Sample-wise log_2_ fold-changes of normalized gene expression with respect to corresponding uninfected samples for genes in Fig. 3A after pseudobulking T cell, NK cell, and ILC2 clusters. P-values of t-test for difference in significance across samples (n=3) marked with * < 0.05, ** < 0.01 or ns: not significant. (**C**) Averaged normalized expression of immunity-related genes for all cells across time colored by hamster species and virus dose. Transparent areas show 95% confidence intervals computed by bootstrap sampling.

### Neutrophils share a common inflammatory state with variable intensity

The strength of the neutrophil response differed drastically between our COVID-19 hamster models and might determine their lethality. To better understand differences in quality of the neutrophil response to SARS-CoV-2, we used a diffusion map approach (*30, 31*) to create a low-dimensional embedding of all neutrophils in Syrian and Roborovski hamsters (Fig. 4A).

**Fig. 4.**
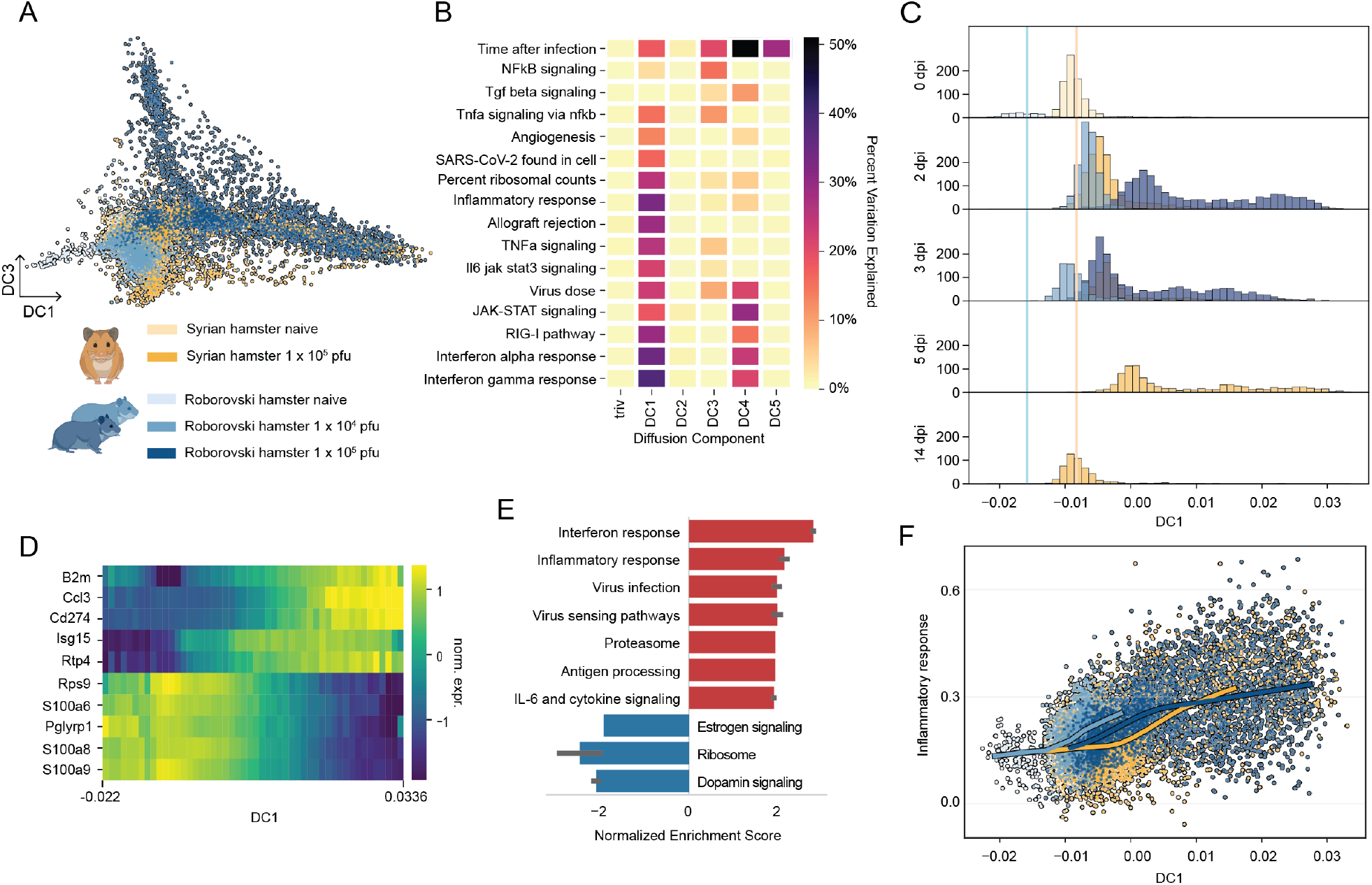
Factor analysis of neutrophils reveals a shared axis of inflammation. (**A**) Low dimensional embedding of first and third diffusion component (DC) for all neutrophils, colored by hamster species and virus dose. (**B**) Percent variance explained by selected covariates for first 5 DCs. (**C**) Distribution of neutrophils along DC1, split by dpi and colored by hamster species and virus dose. Mean position of uninfected cells per hamster type on DC1 is marked with a vertical line. (**D**) Heatmap with gene expression trends for top 5 (anti-)correlating genes with DC1. Gene expression is z-score normalized and convolved by a uniform kernel across seven bins. (**E**) Significant (p-value < 0.05) gene set groups for GSEA on genes ranked by correlation with DC1. Normalized enrichment score expresses the degree of enrichment in genes with higher positive or negative DC1 correlations. (**F**) Inflammatory response score of neutrophils along DC1 colored by hamster species and virus dose. Lines: LOESS regression along DC1.

In this latent space, we identified the first diffusion component (DC1) as the leading variable associated with inflammatory neutrophil responses. This inflammatory axis was characterized by the expression of several inflammatory pathways including JAK-STAT, RIG-I, and TNFα signaling (Fig. 4B). Therefore, we further analyzed this component and visualized the distribution of neutrophils along DC1 for each time point (Fig. 4C). At 0 dpi, neutrophils were localized on the left side of DC1. However, upon infection, there was a general shift to the right side of DC1 for the entire neutrophil population. Specifically, upon high-dose infection, Roborovski hamster neutrophils massively overshot Syrian and low-dose infected Roborovski hamster neutrophils to the far right at 2 dpi. In contrast, Syrian hamster neutrophils progressively moved along DC1, reaching the furthest positions at 5 dpi, before completely returning to the original 0 dpi DC1 “ground state” at 14 dpi.

To identify the molecular drivers that describe DC1, we next ranked genes according to their correlation with the DC1 axis (Fig. 4D, fig. S4A). The left side of DC1 associated with high expression of genes that gradually decreased towards high DC1. These included *S100a8* and *S100a9*, known to compose half of the neutrophils’ cytoplasmic proteins and to act as danger-associate molecular patterns (DAMPs) in inflammation. They critically modulate leukocyte recruitment, cytokine secretion, and neutrophil activation (*32, 33*). Accordingly, the right side of DC1 was marked by increased expression of interferon stimulated and chemokine-encoding genes such as *Isg15, Ccl3*, and *Cd274* (gene encoding for Programmed Death-Ligand 1 (PD-L1)). PD-L1 is expressed by activated neutrophils with lymphocyte suppressive capacities (*11, 34*). Our data show that at 2 and 3 dpi, neutrophils and endothelial cells predominantly express *Cd274* while the expression of the corresponding receptor *Pdcd1* (gene encoding for PD-1) is mainly confined to T and NK cells (fig. S4B). Specifically, expression analysis of *Cd274* in the neutrophil subset confirmed that upregulation is strongest in high-dose infected Roborovski hamsters at 2 dpi (fig. S1C). Accordingly, expression analysis of *Pdcd1* in the T cell subset revealed that the PD-L1 receptor likewise had the highest expression in high-dose infected Roborovski hamsters at that timepoint (fig. S1C). In agreement with these findings, our gene set enrichment analyses (Fig. 4E, fig. S4D) showed that ribosome-specific genes most strongly associated with lower DC1, indicating an initially high and further decreasing ribosomal activity of cells upon infection. In contrast, high DC1 was characterized by the enrichment of multiple pro-inflammatory and immunomodulatory pathways, including TNFα and interferon-mediated signaling. While inflammatory response scores increased with DC1 for all neutrophils (Fig. 4F, fig. S4E), those from high-dose infected Roborovski hamsters extended furthest towards the right side of DC1, indicating an overall higher inflammatory state and lymphocyte suppressive capacity triggered in this species upon infection, matching observations made in severe COVID-19 patients (*11*).

### Identification of a dominant inflammatory response program in endothelial cells

COVID-19 is also a vascular disease (*35*). Nonetheless, endothelial cells, despite their likely importance in determining vascular vulnerability, remain understudied. Due to their limited accessibility, human endothelial cells have primarily been analyzed from post-mortem lung tissues or in vitro systems. To fill this gap, we analyzed endothelial cells analogously to neutrophils. We used a diffusion map approach, followed by gene expression and gene set enrichment analyses to identify the molecular drivers that define characteristic diffusion components (Fig. 5, fig. S5). While the first three DCs described endothelial cell subtype identities (primarily bronchial, vein, artery, and capillary) (Fig. 5A, B), DC4 identified the dominant transcriptional program of endothelial cell responses to SARS-CoV-2 infection, as its variance was explained by a spectrum of pro-inflammatory pathways (Fig. 5B), prompting us to study the dynamics and genes of this component in more detail.

**Fig. 5.**
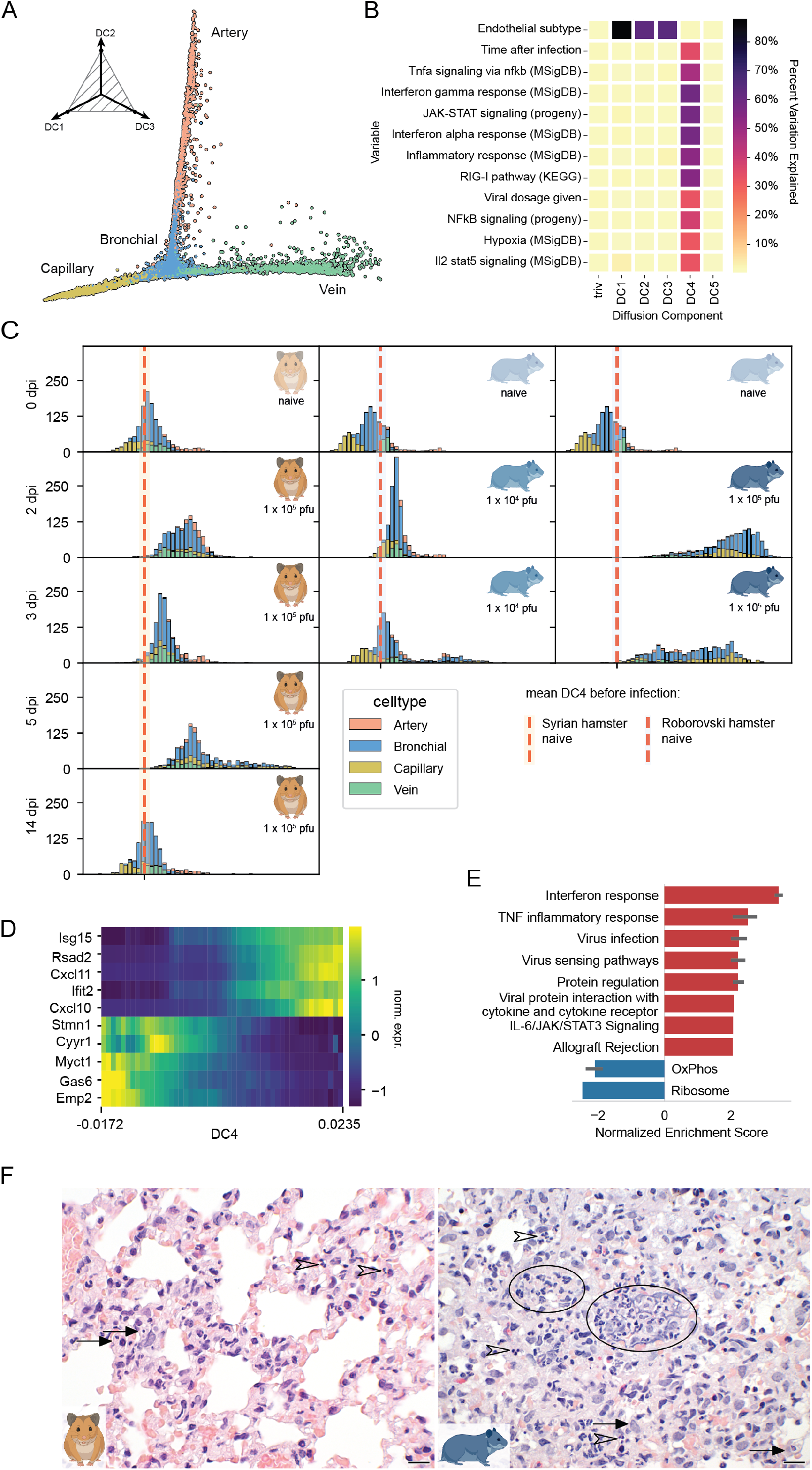
Endothelial cells transiently react to SARS-CoV-2 infection. (**A**) Simplex plot of the first three diffusion components (DCs) for endothelial cells, colored by subtype. (**B**) Variance explained by selected covariates for top five DCs. (**C**) Distribution of cells along DC4, split vertically by hamster species and virus dose and horizontally by dpi, colored by endothelial cell subtype. Average value of DC4 for 0 dpi marked with red vertical line. (**D**) Heatmap with gene expression trends for top 5 (anti-)correlating genes with DC4. Gene expression is z-score normalized and convolved by a uniform kernel across seven bins. (**E**) Significant (p-value < 0.05) gene set groups for GSEA on genes ranked by correlation with DC4. Normalized enrichment score expresses the degree of enrichment in genes with higher positive or negative DC4 correlations. **(F)** Histopathology at 3 days post SARS-CoV-2 infection visualizing reactive neutrophils (open arrowheads) and macrophages (arrows) with higher cell density in the Roborovski hamster. Of note, only Roborovski hamsters developed karyolytic smears / necrotic karyoplasm consistent with alveolar wall and interalveolar blood vessel necrosis (ovals). H&E stain, original magnification = 600x, scale bars = 20 μm.

As seen with neutrophils, endothelial cells were located on the lower / left end of DC4 before infection (0 dpi), however shifted towards the right end of DC4 and took on different DC4 values upon infection (Fig. 5C, fig. S5A). In particular, bronchial and capillary endothelial cells from high-dose infected Roborovski hamsters most rapidly increased in DC4 at 2 dpi before shifting slightly back at 3 dpi. Against this trend, endothelial cells from Syrian hamsters overall only mildly increased in DC4 throughout 2 and 3 dpi, peaking at 5 dpi, and entirely returned to the original state (0 dpi) at 14 dpi. Altogether DC4 is an axis that was shared between hamster species that, upon infection, was occupied much more rapidly and extremely in high-dose infected Roborovski while appearing to be transient in Syrian hamsters. Genes for which the expression correlated positively with increasing DC4 included numerous interferon-stimulated genes such as *Isg15, Rsad2*, and *Ifit2*, as well as chemokines *Cxcl10* and *Cxcl11* (Fig. 5D, fig. S5B), and showed significant upregulation of virus infection / sensing, TNF inflammation, and immune response pathways (Fig. 5E, fig. S5C). For the latter, endothelial cells from high-dose infected Roborovski hamsters scored by far highest in inflammatory response (fig. S5D). While this clearly highlights DC4 as an axis of infection response, genes anti-correlating with progressing DC4 were not classically associated with inflammation, and as already observed for neutrophils, included ribosomal genes, further supporting the hypothesis that translation is actively downregulated for cells while they are strongly reacting to the infectious environment caused by the virus.

Histopathologically, the aforementioned vastly different reactions to the infection in different subtypes of endothelial cells were accompanied by interalveolar wall and blood vessel necrosis with multifocal distinct aggregates of karyolytic smears and cellular debris (Fig. 5F). The larger, primarily venous, endothelial cells from lungs of both hamster species exhibited adhesion and extravasation of leukocytes from the blood as well as dents typical of endotheliitis in both hamster species at 3 dpi. However, interalveolar capillaries and bronchial endothelial cells, while mildly affected in Syrian hamsters, were strongly distressed by necrosis in Roborovski hamsters at 2 and 3 dpi, especially in the higher dose group. These different reactions by endothelial subtype are also visible on DC4, where capillary endothelial cells, particularly from Roborovski hamsters, are much further on the right compared to arterial and venous subtypes.

## Discussion

Since the onset of the SARS-CoV-2 pandemic, a vast amount of knowledge has been gathered on the pathogenesis of COVID-19, the disease caused by this virus. A variety of animal models and patient cohort data revealed details on mechanisms of pathogenesis (*14, 36*), immunity (*37, 38*), and response to therapy (*39*), which greatly improved our understanding of the disease. Macroscopic risk factors for severe disease such as age and specific comorbidities have been identified (*40*), and also some molecular aspects such as interferon autoantibodies or HLA genotype (*41-43*). Still, the fundamental question of the initial cellular mechanisms that direct the path to mild, moderate, or severe disease remains unsolved. There are two key obstacles to answering this question using human lung patient data. Firstly, such data are not available for early time points post infection. Secondly, bronchoalveolar lavages, which represent the typical sampling method, cover soluble immune cells well, but tissue cells only to a very limited extent.

In order to overcome these hurdles, i.e. to comparatively study moderate and severe disease at its very onset, we here probed published and previously unpublished scRNA-seq datasets from patients and two COVID-19 hamster models for correlates of disease severity. Syrian hamsters show a moderate course of COVID-19 with subsequent recovery, whereas Roborovski dwarf hamsters frequently succumb to severe disease (*18*). scRNA-seq data allow for unbiased, transcriptome-wide investigations of cellular activation states at single-cell resolution. It can therefore provide a comprehensive overview of the cellular processes happening in COVID-19. By combining this and sampling of animals with different disease severity, we aimed to investigate early processes in the disease that may lead to different outcomes.

In our analysis, Syrian hamsters displayed an efficient type 1 immune response, including NK cell and monocyte recruitment into lungs. This is in line with patient data describing that a moderate course of COVID-19 correlates with functional NK cells (*44*). The same study found that augmented expression levels of transforming growth factor-β (TGFβ) suppressed NK cells in their anti-viral properties, fostering a severe disease course. Following their efficient, initial innate response, Syrian hamsters generally survived beyond 3 dpi and mounted effective T cell and neutralizing antibody responses against SARS-CoV-2 (*19*), which was not observed in Roborovski hamsters. This could mean that this severe disease animal model does not reach this state of efficient viral clearance early enough, or that the inability to initiate this response is one root cause for severe disease.

In contrast to Syrian hamsters, in which we observed clear upregulation of type 1 lineage-determining transcription factors *Tbx21* and *Eomes*, we did not observe upregulation of the type 3 lineage-determining transcription factor *Rorc* in Roborovski hamsters. Similarly, *Stat3* a transcription factor in the IL-6, IL-21 and IL-23 signaling pathways for Th17 cells, was upregulated in Syrian but not Roborovski hamsters. However, presence of cytokines *Il6* and *Il23a*, as well as *Socs1*, indicates, together with the absence of type 1 and type 2 associated transcription patterns, a type 3 response in Roborovski hamsters with severe disease course.

The cell population most strikingly related to severity of inflammation and disease in patients, as well as hamsters, are neutrophils. Neutrophils are the downstream effector cells of type 3 immunity, providing defense against extracellular pathogens, such as bacteria and fungi at epithelial barriers (*25*). However, neutrophils are also associated with autoimmune disease and infection-triggered immunopathology (*25, 45-48*). The protective role of neutrophils and type 3 immunity against viral infections is limited to specific cases, e.g. HSV-1 infection of the cornea, rhinovirus infection, and influenza infection (*49-51*). Importantly, in the context of SARS-CoV-2 infection, Th17 cells and neutrophils have been associated with increased disease severity and immunopathology (*11, 52*). Using diffusion analysis, we found a latent factor in neutrophils uniquely characterizing their response to the infection across hamster species, which revealed an overly exacerbated neutrophil response in Roborovski hamsters. Based on this finding, our data clearly link inflammatory neutrophils to disease severity and aligns with previous studies in which we showed that dampening neutrophilic inflammation by dexamethasone treatment prevents lethal disease outcomes (*53*). Moreover, inflammatory neutrophils in severely diseased Roborovski hamsters strongly expressed *Cd274*, as do human neutrophils in patients with a severe disease course (*52*). However, the origin of the neutrophil response remains obscure.

An important hallmark of COVID-19 pathogenesis is endothelial damage across multiple organs. Since endothelial cells are challenging to study in human patients, animal models may be of particular use. In our diffusion analysis, we identified an axis of inflammation (DC4), i.e. a specific gene expression program marked by upregulation of pro-inflammatory genes that endothelial cells follow upon infection. This effect was particularly strong for bronchial and capillary cells of high-dose infected Roborovski hamsters. In the histopathological analysis, we observed endothelial cell necrosis, particularly in high-dose infected Roborovski hamsters. Still, it is important to note that endothelial cells were also strongly activated along DC4 in Syrian hamsters, but reverted to ground state, and did not show obvious histological endothelial damage. It thus appears that endothelial activation is associated with disease severity, but not necessarily causative, and under beneficial conditions also reversible.

Our data from lung tissues offer a unique perspective on the involvement of endothelial cells in the pathological development of a severe disease course, given the lack of corresponding data and samples from humans. Notably, our data suggest that immunological events early in the course of SARS-CoV-2 infection cause development of either type 1 or type 3-biased immune responses, thereby determining disease progression in both hamsters and humans. Future research can make use of the Roborovski hamster model for severe COVID-19-like disease to further elucidate mechanisms of severe disease and investigate potential medical interventions.

## Acknowledgements

We thank Diether Lambrechts and Roland Eils for sharing raw sequencing data not used in this study. Hamster and human icons were created with BioRender.com. Computation was in part performed on the HPC for Research cluster of the Berlin Institute of Health.

## Author contributions

Conceptualization: SP, GN, EW, MW, JT, ML, SDP

Methodology: SP, GN, PP, EW, JT, SDP

Investigation: SP, GN, EW, JMA, SK, AV, JK, FP, PP, DP, LGTA, ADG, JT, ML, SDP

Formal analysis: SP, SDP

Visualization: SP, GN, SK, AV, ADG, SDP

P Project administration: GN, JT, SDP

Supervision: GN, CG, ADG, NB, MW, JT, ML, SDP

Writing – original draft: SP, GN, EW, JT, SDP

Writing – review & editing: SP, GN, EW, SK, PP, ADG, JT, ML, SDP

## Data and materials availability

Count data are available from the corresponding author upon reasonable request. Code for data analyses is available at GitHub (https://github.com/stefanpeidli/PanCov19_Hamster).

## Supplementary Materials

Materials and Methods

Figs. S1 to S5

## References

1. J. E. Michalski, J. S. Kurche, D. A. Schwartz, From ARDS to pulmonary fibrosis: the next phase of the COVID-19 pandemic? Transl Res 241, 13–24 (2022).

2. A. Mohammadi et al., Post-COVID-19 Pulmonary Fibrosis. Cureus 14, e22770 (2022).

3. S. A. Jasim et al., The deciphering of the immune cells and marker signature in COVID-19 pathogenesis: An update. J Med Virol 94, 5128–5148 (2022).

4. R. Knoll, J. L. Schultze, J. Schulte-Schrepping, Monocytes and Macrophages in COVID-19. Front Immunol 12, 720109 (2021).

5. S. M. Vora, J. Lieberman, H. Wu, Inflammasome activation at the crux of severe COVID-19. Nature Reviews Immunology 21, 694–703 (2021).

6. F. V. S. Castanheira, P. Kubes, Neutrophils during SARS-CoV-2 infection: Friend or foe? Immunol Rev 314, 399–412 (2023).

7. S. R. Paludan, T. H. Mogensen, Innate immunological pathways in COVID-19 pathogenesis. Science Immunology 7, eabm5505 (2022).

8. R. A. Hernandez Acosta, Z. Esquer Garrigos, J. R. Marcelin, P. Vijayvargiya, COVID-19 Pathogenesis and Clinical Manifestations. Infect Dis Clin North Am 36, 231–249 (2022).

9. K. E. Remy et al., Severe immunosuppression and not a cytokine storm characterizes COVID-19 infections. JCI Insight 5, (2020).

10. D. Wendisch et al., SARS-CoV-2 infection triggers profibrotic macrophage responses and lung fibrosis. Cell 184, 6243–6261.e6227 (2021).

11. J. Schulte-Schrepping et al., Severe COVID-19 Is Marked by a Dysregulated Myeloid Cell Compartment. Cell 182, 1419–1440 e1423 (2020).

12. E. Falcinelli, E. Petito, P. Gresele, The role of platelets, neutrophils and endothelium in COVID-19 infection. Expert Rev Hematol 15, 727–745 (2022).

13. R. Zhang et al., Neutrophil autophagy and NETosis in COVID-19: perspectives. Autophagy, 1–10 (2022).

14. C. Fan et al., Animal models for COVID-19: advances, gaps and perspectives. Signal Transduct Target Ther 7, 220 (2022).

15. F. Qi, C. Qin, Characteristics of animal models for COVID-19. Animal Model Exp Med 5, 401–409 (2022).

16. A. D. Gruber, T. C. Firsching, J. Trimpert, K. Dietert, Hamster models of COVID-19 pneumonia reviewed: How human can they be? Veterinary Pathology 59, 528–545 (2022).

17. H. Chu, J. F. Chan, K. Y. Yuen, Animal models in SARS-CoV-2 research. Nat Methods 19, 392–394 (2022).

18. J. Trimpert et al., The Roborovski Dwarf Hamster Is A Highly Susceptible Model for a Rapid and Fatal Course of SARS-CoV-2 Infection. Cell Rep 33, 108488 (2020).

19. G. Nouailles et al., Temporal omics analysis in Syrian hamsters unravel cellular effector responses to moderate COVID-19. Nat Commun 12, 4869 (2021).

20. R. L. Chua et al., COVID-19 severity correlates with airway epithelium–immune cell interactions identified by single-cell analysis. Nature Biotechnology 38, 970–979 (2020).

21. M. Liao et al., Single-cell landscape of bronchoalveolar immune cells in patients with COVID-19. Nature Medicine 26, 842–844 (2020).

22. J. C. Melms et al., A molecular single-cell lung atlas of lethal COVID-19. Nature 595, 114–119 (2021).

23. N. Osterrieder et al., Age-Dependent Progression of SARS-CoV-2 Infection in Syrian Hamsters. Viruses 12, 779 (2020).

24. Y. J. Hou et al., SARS-CoV-2 Reverse Genetics Reveals a Variable Infection Gradient in the Respiratory Tract. Cell 182, 429–446.e414 (2020).

25. F. Annunziato, C. Romagnani, S. Romagnani, The 3 major types of innate and adaptive cell-mediated effector immunity. Journal of Allergy and Clinical Immunology 135, 626–635 (2015).

26. J. Zhang et al., Sequential actions of EOMES and T-BET promote stepwise maturation of natural killer cells. Nat Commun 12, 5446 (2021).

27. L. Zhou, Striking similarity: GATA-3 regulates ILC2 and Th2 cells. Immunity 37, 589–591 (2012).

28. H. Song et al., Critical role for GATA3 in mediating Tie2 expression and function in large vessel endothelial cells. J Biol Chem 284, 29109–29124 (2009).

29. M. L. Sobah, C. Liongue, A. C. Ward, SOCS Proteins in Immunity, Inflammatory Diseases, and Immune-Related Cancer. Front Med (Lausanne) 8, 727987 (2021).

30. R. R. Coifman, S. Lafon, Diffusion maps. Applied and Computational Harmonic Analysis 21, 5–30 (2006).

31. L. Haghverdi, F. Buettner, F. J. Theis, Diffusion maps for high-dimensional single-cell analysis of differentiation data. Bioinformatics 31, 2989–2998 (2015).

32. S. Wang et al., S100A8/A9 in Inflammation. Front Immunol 9, 1298 (2018).

33. E. G. G. Sprenkeler et al., S100A8/A9 Is a Marker for the Release of Neutrophil Extracellular Traps and Induces Neutrophil Activation. Cells 11, (2022).

34. S. de Kleijn et al., IFN-γ-stimulated neutrophils suppress lymphocyte proliferation through expression of PD-L1. PLoS One 8, e72249 (2013).

35. H. K. Siddiqi, P. Libby, P. M. Ridker, COVID-19 - A vascular disease. Trends Cardiovasc Med 31, 1–5 (2021).

36. S. Choudhary, I. Kanevsky, L. Tomlinson, Animal models for studying COVID-19, prevention, and therapy: Pathology and disease phenotypes. Vet Pathol 59, 516–527 (2022).

37. C. Brady, T. Tipton, S. Longet, M. W. Carroll, Pre-clinical models to define correlates of protection for SARS-CoV-2. Front Immunol 14, 1166664 (2023).

38. S. Clever, A. Volz, Mouse models in COVID-19 research: analyzing the adaptive immune response. Med Microbiol Immunol 212, 165–183 (2023).

39. G. Li, R. Hilgenfeld, R. Whitley, E. De Clercq, Therapeutic strategies for COVID-19: progress and lessons learned. Nat Rev Drug Discov 22, 449–475 (2023).

40. Y. D. Gao et al., Risk factors for severe and critically ill COVID-19 patients: A review. Allergy 76, 428–455 (2021).

41. Q. Zhang, P. Bastard, A. Cobat, J. L. Casanova, Human genetic and immunological determinants of critical COVID-19 pneumonia. Nature 603, 587–598 (2022).

42. J. V. Garmendia, A. H. García, C. V. De Sanctis, M. Hajdúch, J. B. De Sanctis, Autoimmunity and Immunodeficiency in Severe SARS-CoV-2 Infection and Prolonged COVID-19. Curr Issues Mol Biol 45, 33–50 (2022).

43. D. G. Augusto et al., A common allele of HLA is associated with asymptomatic SARS-CoV-2 infection. Nature, (2023).

44. M. Witkowski et al., Untimely TGFβ responses in COVID-19 limit antiviral functions of NK cells. Nature 600, 295–301 (2021).

45. M. M. Saleh, W. A. Petri, Jr., Type 3 Immunity during Clostridioides difficile Infection: Too Much of a Good Thing? Infect Immun 88, (2019).

46. L. Borkner, L. M. Curham, M. M. Wilk, B. Moran, K. H. G. Mills, IL-17 mediates protective immunity against nasal infection with Bordetella pertussis by mobilizing neutrophils, especially Siglec-F(+) neutrophils. Mucosal Immunol 14, 1183–1202 (2021).

47. Y. J. Lu et al., Interleukin-17A mediates acquired immunity to pneumococcal colonization. PLoS Pathog 4, e1000159 (2008).

48. Y. Zheng et al., Interleukin-22, a T(H)17 cytokine, mediates IL-23-induced dermal inflammation and acanthosis. Nature 445, 648–651 (2007).

49. B. Kim, P. P. Sarangi, A. K. Azkur, S. D. Kaistha, B. T. Rouse, Enhanced viral immunoinflammatory lesions in mice lacking IL-23 responses. Microbes Infect 10, 302–312 (2008).

50. S. Wiehler, D. Proud, Interleukin-17A modulates human airway epithelial responses to human rhinovirus infection. Am J Physiol Lung Cell Mol Physiol 293, L505–515 (2007).

51. H. Hamada et al., Tc17, a unique subset of CD8 T cells that can protect against lethal influenza challenge. J Immunol 182, 3469–3481 (2009).

52. M. Orlov, P. L. Wander, E. D. Morrell, C. Mikacenic, M. M. Wurfel, A Case for Targeting Th17 Cells and IL-17A in SARS-CoV-2 Infections. J Immunol 205, 892–898 (2020).

53. E. Wyler et al., Key benefits of dexamethasone and antibody treatment in COVID-19 hamster models revealed by single-cell transcriptomics. Mol Ther 30, 1952–1965 (2022).

54. M. Kanehisa, S. Goto, KEGG: kyoto encyclopedia of genes and genomes. Nucleic Acids Res 28, 27–30 (2000).

55. M. Kanehisa, Toward understanding the origin and evolution of cellular organisms. Protein Sci 28, 1947–1951 (2019).

56. M. Kanehisa, M. Furumichi, Y. Sato, M. Kawashima, M. Ishiguro-Watanabe, KEGG for taxonomy-based analysis of pathways and genomes. Nucleic Acids Res 51, D587–d592 (2023).

57. A. Liberzon et al., The Molecular Signatures Database (MSigDB) hallmark gene set collection. Cell Syst 1, 417–425 (2015).

58. A. Subramanian et al., Gene set enrichment analysis: A knowledge-based approach for interpreting genome-wide expression profiles. Proceedings of the National Academy of Sciences 102, 15545–15550 (2005).

59. R. Wölfel et al., Virological assessment of hospitalized patients with COVID-2019. Nature 581, 465–469 (2020).

60. J. Trimpert et al., Live attenuated virus vaccine protects against SARS-CoV-2 variants of concern B.1.1.7 (Alpha) and B.1.351 (Beta). Sci Adv 7, eabk0172 (2021).

61. J. M. Adler et al., A non-transmissible live attenuated SARS-CoV-2 vaccine. Mol Ther, (2023).

62. J. Trimpert et al., Development of safe and highly protective live-attenuated SARS-CoV-2 vaccine candidates by genome recoding. Cell Rep 36, 109493 (2021).

63. F. A. Wolf, P. Angerer, F. J. Theis, SCANPY: large-scale single-cell gene expression data analysis. Genome Biology 19, (2018).

64. M. I. Love, W. Huber, S. Anders, Moderated estimation of fold change and dispersion for RNA-seq data with DESeq2. Genome Biology 15, (2014).

65. R. Lopez, J. Regier, M. B. Cole, M. I. Jordan, N. Yosef, Deep generative modeling for single-cell transcriptomics. Nature Methods 15, 1053–1058 (2018).

66. F. Mölder et al., Sustainable data analysis with Snakemake. F1000Research 10, 33 (2021).

67. S. Andreotti et al., De Novo-Whole Genome Assembly of the Roborovski Dwarf Hamster (Phodopus roborovskii) Genome: An Animal Model for Severe/Critical COVID-19. Genome Biol Evol 14, (2022).

68. S. L. Wolock, R. Lopez, A. M. Klein, Scrublet: Computational Identification of Cell Doublets in Single-Cell Transcriptomic Data. Cell Systems 8, 281–291.e289 (2019).

69. L. McInnes, J. Healy, J. Melville, UMAP: Uniform Manifold Approximation and Projection for Dimension Reduction. arXiv pre-print server, (2020).

70. R. Satija, J. A. Farrell, D. Gennert, A. F. Schier, A. Regev, Spatial reconstruction of single-cell gene expression data. Nat Biotechnol 33, 495–502 (2015).

71. C. Hafemeister, R. Satija, Normalization and variance stabilization of single-cell RNA-seq data using regularized negative binomial regression. Genome Biol 20, 296 (2019).

72. M. I. Love. (2023), vol. 2023.

73. D. Smedley et al., BioMart – biological queries made easy. BMC Genomics 10, 22 (2009).

74. T. Stuart et al., Comprehensive Integration of Single-Cell Data. Cell 177, 1888–1902.e1821 (2019).

75. Z. Fang, X. Liu, G. Peltz, GSEApy: a comprehensive package for performing gene set enrichment analysis in Python. Bioinformatics 39, (2023).

