## Supplementary Materials for "Single-cell-resolved interspecies comparison identifies a shared inflammatory axis and a dominant neutrophil-endothelial program in severe COVID-19"

**The PDF file includes:**

Materials and Methods
Figs. S1 to S5

### Materials and Methods

#### Animal Husbandry

The animal experiments were approved by the competent state authority (Landesamt für Gesundheit und Soziales Berlin, Germany, approval number 0086/20) and were performed in accordance with national and international regulations. In accordance with the 3R principle, no additional animal experiments were performed for this study. Instead, we used data and samples from animals studied in our previously published research (*18, 19, 23*). In these experiments, female and male Syrian hamsters (Mesocricetus auratus; RjHan:AURA, Janvier Labs, Saint-Berthevin, France) and Roborovski hamsters (Phodopus roborovskii, German pet trade) were housed in a BSL-3 facility in individually ventilated cages (IVCs; Tecniplast, Buguggiate, Italy) with abundant enrichment (Carfil, Oud-Turnhout, Belgium) and ad libidum access to food and water. Cage temperature and relative humidity were recorded daily and ranged from 22 to 24 °C and 40 to 55%, respectively. Animals were allowed at least 7 days to acclimatize before the start of the experiments.

#### Cells and Virus

The SARS-CoV-2 isolate (BetaCoV/Germany/BavPat1/2020) (*59*) was kindly provided by Drs. Daniela Niemeyer and Christian Drosten, Charité Berlin, Germany. Virus stocks for animal experiments were obtained by propagating the virus under BSL-3 conditions on Vero E6 cells (ATCC CRL-1586) in minimal essential medium (MEM; PAN Biotech, Aidenbach, Germany) supplemented with 10% fetal bovine serum (FBS; PAN Biotech, Aidenbach, Germany), 100 IU/ml penicillin G, and 100 µg/ml streptomycin (Carl Roth, Karlsruhe, Germany). Prior to animal testing, low-passage stocks were titrated to Vero E6 cells using semi-solid overlay medium as described (*60*). Briefly, Vero E6 cells were incubated with serial 10-fold dilutions of virus strains for 2 hours. The virus inoculum was then replaced with an overlay medium consisting of Dulbecco's modified Eagle's medium (DMEM; PAN Biotech, Aidenbach, Germany), 2.5% microcrystalline cellulose (Avicel RC-591; DuPont, Wilmington, DE, USA) and 10% FBS. After 72 hours of incubation at 37°C in a 5% $CO_2$ atmosphere, cells were fixed with 4% formaldehyde for 24 hours and plaques were visualized by methylene blue counterstaining. Sequence integrity of virus stocks was determined by Illumina sequencing as described (*61*) and aligned to the isolate reference sequence (GenBank: MT270101 and GISAID: EPI_ISL_406862).

#### Animal Experimentation

10- to 12-week-old male and female Syrian and Roborovski hamsters were infected intranasally with $1x10^5$ or $1x10^4$ plaque-forming units (pfu) SARS-CoV-2 (variant B1, isolate BetaCoV/Germany/BavPat1/2020) under anesthesia as previously described (*62*). To prevent any prolonged suffering, the hamsters were clinically examined twice daily. Animals with body weight loss >15% for more than 48 hours were euthanized according to the animal use protocol. Otherwise, naive hamsters (n = 3) and hamsters at 2, 3, 5, and 14 days post infection (n = 3 each) were randomly selected. Euthanasia was performed by cervical dislocation and exsanguination under anesthesia as previously described (*23*). Among other materials, all lung lobes were collected for subsequent analyses. Specifically, the left lobe was used for histopathology, the right caudal lobe for single-cell analysis, the right cranial lobe for virological measurements, and the right middle lobe for bulk RNA and proteomic analysis as described (*19*).

**Single-Cell Isolation**

Established cell isolation protocols were modified to comply with BSL3 facility regulations. For single cell isolation, the caudal lobes of the right lung were stored in 1× PBS, 0.5% BSA containing 2 µg/ml actinomycin D. The lobes were dissociated mechanically and enzymatically. Tissue was first disrupted with forceps for 2 minutes in specific digestion medium (3.4 mg/mL collagenase Cls II (Merck), 1 mg/mL DNase I (PanReac AppliChem) in 2 mL Dispase medium per lung lobe (Corning), 50 caseinolytic units/mL) and then enzymatically digested at 37°C and 5% $CO_2$ for 30 minutes. The cell suspension was further dissociated by pipetting, and filtered through 70 µm cell strainers. The suspensions were centrifuged at 350 g for 6 minutes at 4 °C, and the pellets were subjected to erythrocyte lysis by resuspension in appropriate buffer (BioLegend). The reaction was stopped by washing with PBS/BSA buffer and the cells were centrifuged. Cells were resuspended in low BSA buffer (1 × PBS, 0.04% BSA) and then filtered through 40 µm FloMi filters (Merck). Cell number and viability were determined microscopically using trypan blue.

**Generation of Single-Cell RNA-Sequencing Data**

Sequencing libraries were generated using the 3’ Chromium Next GEM Single Cell 3’ Reagent kit (10x Genomics) according to the manufacturer’s instructions, and sequenced on a NovaSeq 6000 device (Illumina) to a depth of about 300 million reads per sample.

**Histopathology**

For histopathological evaluation, left lungs were immersion fixed in 10 % buffered formalin (pH 7.0) for 48 h, processed overnight in a Tissue Tek robot, embedded in paraffin and cut at 2 µm thickness. Sections were stained with hematoxylin and eosin (HE). Photomicrographs were taken with an Olympus BX41 microscope, a DP80 Microscope Digital Camera (Olympus, Hamburg, Germany), and cellSensTM Imaging Software, Version 1.18 (Olympus Soft Imaging Solutions).

**Data Analysis**

Data analysis was primarily performed using scanpy (*63*) (v1.9.1) and DESeq2 (*64*) (v1.38.3). For integration we used scVI (*65*) (scvi-tools, v0.19.0). We relied on snakemake

(*66*) (v7.20.0) to compose a reproducible and modular analysis pipeline using parallelized computations. Code used to analyze the data, produce the figures and accompanying source tables in the manuscript is publicly available at https://github.com/stefanpeidli/pancov19_singlecell_comparison.

##### *Processing of single-cell RNA-seq data*

Raw scRNA-seq reads (FASTQ format) were aligned to reference genomes [Roborovski hamster: (*67*) and Syrian hamster: MesAur1.0 assembly for Mesocricetus auratus from ENSEMBL] and quantified to produce a count matrix using CellRanger (v6.1.1) from 10X Genomics with default parameters. Filtered feature barcode matrices from cellranger count were read into scanpy. Raw counts were saved to 'adata.layers['counts']' for later use. We normalized counts using 'scanpy.pp.normalize_total' and log-scaled the results using 'scanpy.pp.log1p'. We did not z-score normalize the gene expression matrix. Doublets were identified and removed using scrublet (v0.2.3) (*68*). We used UMAP (*69*) for visual embeddings only, and diffusionmap (*30, 31*) to generate latent spaces for further analysis. Gene signatures were called using scanpy.tl.score_genes.

##### *Filtering, clustering, and cell type annotation*

Following generation of count matrices using CellRanger, the data was further processed in R using Seurat (*70*). For both hamster species, all samples were merged into one Seurat object, and cells grouped into 45-50 clusters. Based on known marker genes, clusters were annotated into broader cell type categories, or "mixed/unknown". Subsequently, clusters representing likely low quality cells, i.e. low UMI and gene counts, were discarded. In all other clusters, cells with less than the median of UMI counts, or more than the upper quartile plus three inter-quartile ranges (outliers) were discarded. In contrast to a general quality threshold based on UMI or gene counts, this introduces less bias as e.g. neutrophils have much lower UMI/gene counts than macrophages. After that first step, samples were integrated using the sctransform workflow (*71*), and clustered again, to about 30 clusters. As before, these clusters were assigned to cell types using marker genes.

##### *Differential expression analysis*

We averaged unnormalized counts per sample and cell type in python and exported the resulting pseudo bulk count matrix to be used in the DESeq2 package (*64*) (v1.38.3) in R. We filtered out genes with less than 10 counts in total, and pseudo bulks obtained by summing over less than 10 cells. Adhering to recommendations by DESeq2 authors on applications to scRNA-seq data (*72*), we used the following options: test="LRT", minReplicatesForReplace=Inf. We compared all infected samples (2 and 3 dpi only for hamsters) against control / healthy samples separately for each hamster type and human dataset. DESeq2 results were then exported and analyzed in python, where we generally worked with results filtered by adjusted p-value of 0.05 if not otherwise noted.

##### *Mapping orthologous genes between hamster and human genomes*

Instead of "upper-casing" hamster genes to get human orthologs we used existing orthologies to map genes more accurately. We exported human and mouse orthologs from biomart (*73*), since the hamster genomes used in this study were originally aligned against mouse genomes. For each hamster gene, we searched whether there were human orthologs in the database. If multiple were found, we used the best match according to orthology confidence reported by biomart. If no ortholog was found, we kept the gene name and upper-cased it as a best-guess matching.

##### *Integration of hamster data with scVI*

After concatenating all Syrian and Roborovski hamster scRNA-seq data, we identified the 2000 most highly variable genes using 'scanpy.pp.highly_variable_genes' on unnormalized counts using flavor='seurat_v3' (*74*), giving organism identity (Syrian/Roborovski) as batch_key. This selects for genes that are highly variable within Syrian and Roborovski and can be seen as a mild form of batch effect correction. We ran scVI (*65*) with hamster type as batch_key without further covariate keys until convergence (early_stopping=True), with training/test/validation set size ratios of 90/5/5 %, respectively. Integrations were both performed on all data and on subsets only (macrophages / endothelial cells / neutrophils) to achieve appropriate resolution in the scVI latent spaces in each case. When not otherwise mentioned, the scVI latent spaces were used for computing KNN graphs and therefore also for the KNN-based UMAP and diffusion map embeddings.

##### *Post-hoc interpretation of diffusion components*

We employed a simple strategy for post-hoc interpretation of latent variables using one-way ANOVA: Given a continuous or categorical explanatory variable per cell, for each latent variable we fit a simple linear regression model. We then computed how much variability between cells across the latent variable can be explained by the explanatory variable in this linear model as:

$$Variation\ Explained = 1 - \frac{RSS}{TSS} \times DOF_{adjustment}$$

with variables RSS: residual sum of squares, TSS: total sum of squares, and $DOF_{adjustment}$: degrees of freedom adjustment factor. These values were obtained from a simple uni-variate linear regression against each latent factor separately. As explanatory variables we used variables that we would expect to play a significant role in shaping the data by prior knowledge, e.g. cell cycle phases, virus detected, cell subtype, various signaling pathway scores, or study variables such as days past infection, organism or virus dose. In essence, the resulting variation explained describes how much variation in the data can be explained by a linear model based on the current predictor such as cell subtype or cell cycle phase. We applied this post-hoc interpretation method to the latent variables produced by PCA of mean-shifted 2000 highly-variable genes (HVG) expression, scVI, and diffusion map on scVI. Among those, diffusion map resulted in high overall interpretability with latent variables explained by few distinct explanatory variables, whereas both PCA and scVI tended to have few latent dimensions explained by many overlapping explanatory variables. In short, diffusion map model distances between cells as a diffusion process (random walk) on the data manifold. The resulting diffusion components describe various aspects of the data in a continuous way and are naturally ordered by "relevance" by virtue of being defined by an eigendecomposition of the diffusion matrix defined on the KNN graph of the data.

##### *Gene set enrichment analysis*

For gene set enrichment analysis (GSEA) (*58*) we utilized the python package gseapy (*75*) (v0.10.8). For comparison between infected and control samples, results from DESeq2 (infected vs. control / healthy) were first filtered to exclude genes with less than 10 baseMean. The prerank module by gseapy takes in a ranked list of genes together with the score used for ranking them. As basis for the score we used negative log10 of p-values prior to multiple-testing correction, as this correction does not change the ranking except that it might lead to genes having ties in scores which should generally be avoided in GSEA. Subsequently, these values, representing the statistical strength of each gene's change, were multiplied by the sign of each gene's log fold change, representing the direction of change. This score was used to rank genes. For GSEA of neutrophils and endothelial cells, we ranked all genes by spearman correlation with the corresponding diffusion component. Afterwards, we selected genes with a higher number of counts over all cells than the median to be considered. In both cases, ranked genes were given to gseapy.prerank, with parameters: min_size=15, max_size=1000, permutation_num=1000. Gene sets used comprise KEGG_2021_Human (*54*) and MSigDB_Hallmark_2020 (*57*). GSEA results with an FDR q-value < 0.05 were used for further analysis, including the normalized enrichment scores as direct output from gseapy.

### Supplementary Figures

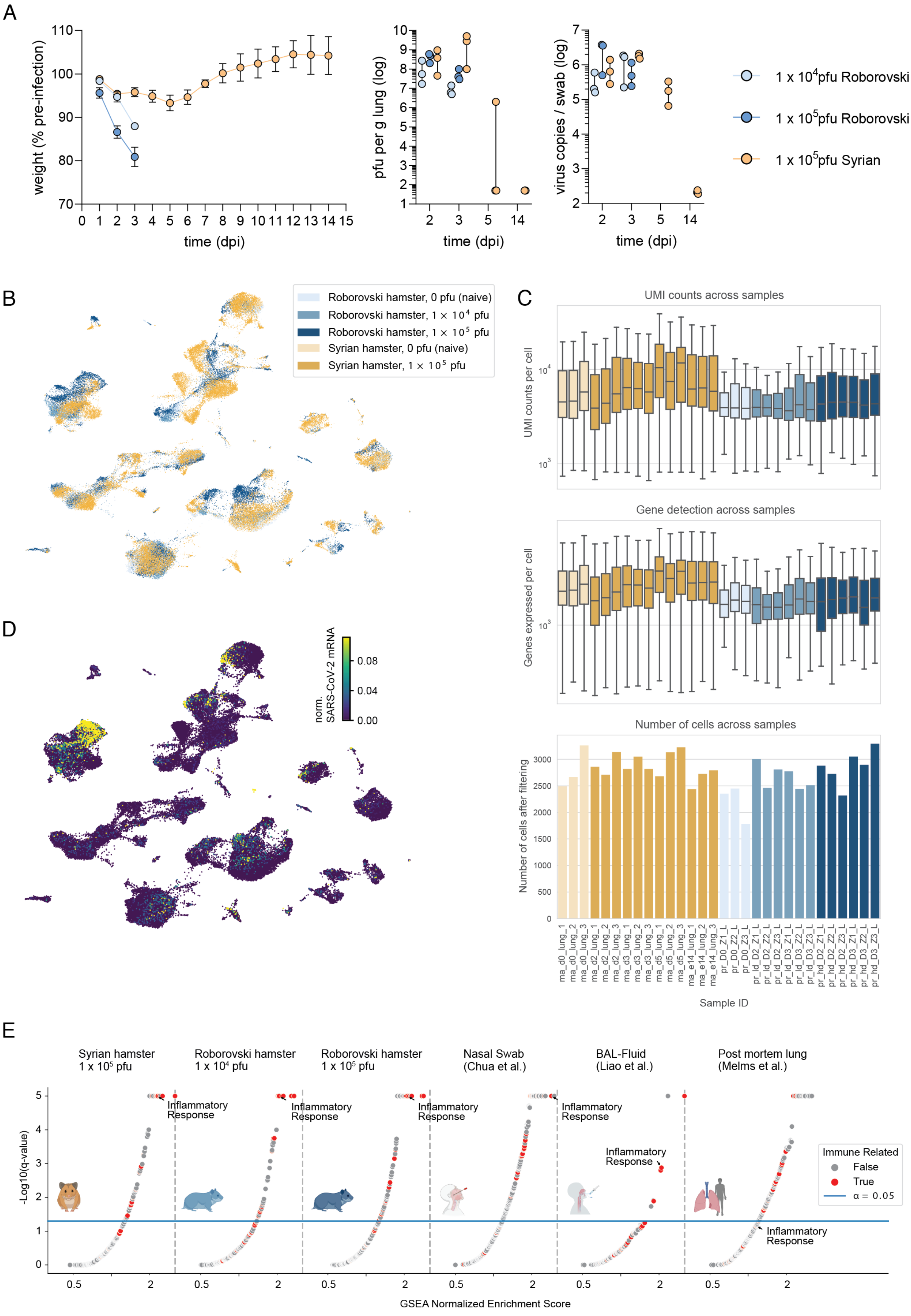

**fig. S1 Clinical data, scRNA-seq data statistics, upregulated gene sets across datasets.** (**A**) Weight w.r.t. pre-infected (left), log-scale virus pfu per gram lung tissue (middle), and log-scale virus copies per swab (right) colored by study branch (hamster type and virus dose) along dpi (clinical data previously published: (*18, 19*). (**B**) UMAP embedding of integrated datasets colored by hamster species and virus dose. (**C**) UMI counts (top), number of genes with at least one UMI count (middle) per cell and number of cells after filtering (bottom) per sample colored by hamster species. (**D**) UMAP embedding as in B showing normalized SARS-CoV-2 mRNA sequence counts per cell. (**E**) GSEA results sets across datasets based on DEGs from DESeq2 testing infected (2 and 3 dpi) vs. control showing upregulated gene sets with immune-related terms indicated in red. Normalized Enrichment Score (NES) as x-axis and -log10(q-value) as y-axis. Blue line corresponds to a q-value of 0.05.

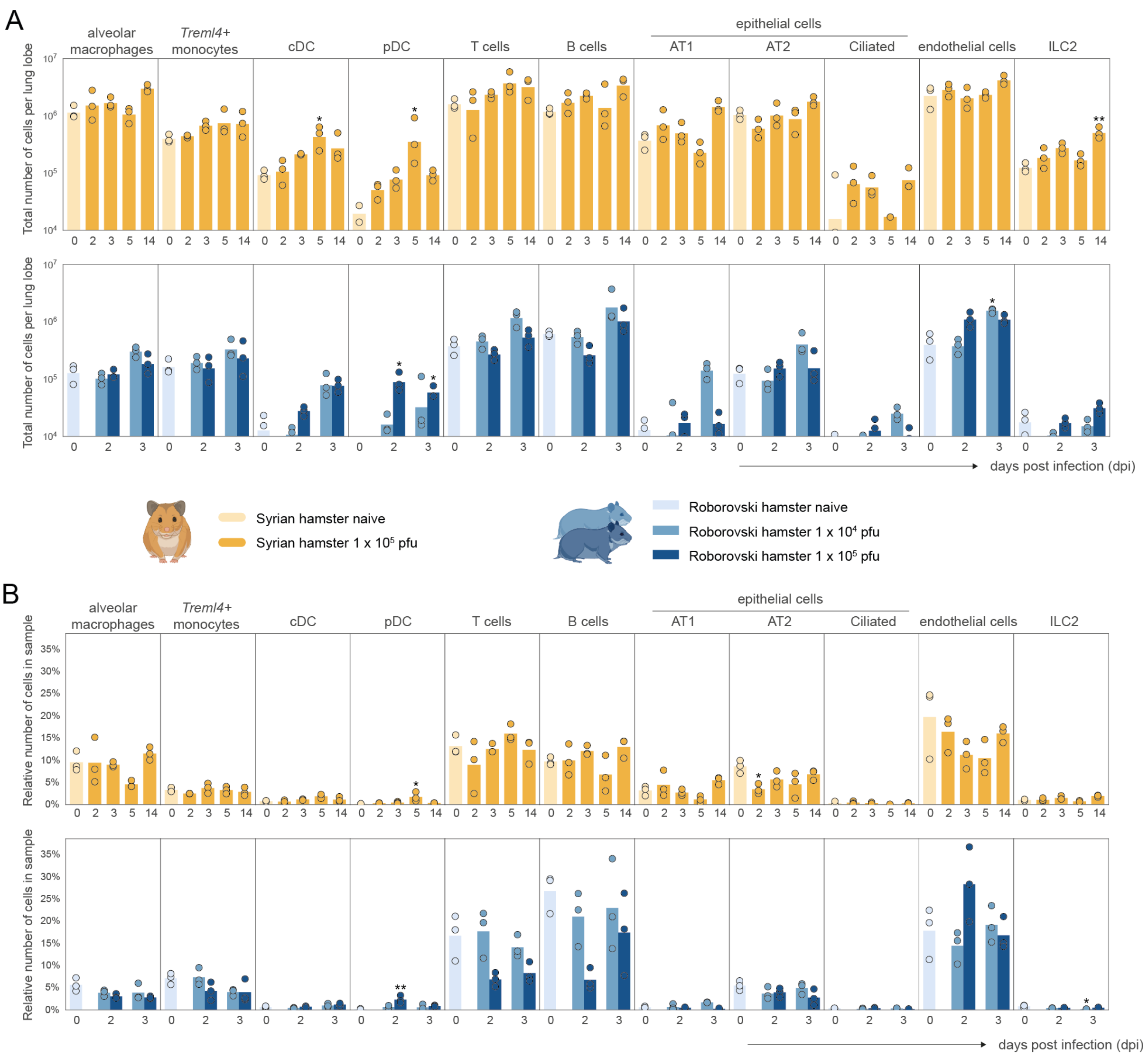


**fig. S2 Cell numbers across all other annotated cell types**. (**A**) as in Fig. 2B. (**B**) as in Fig. 2C.

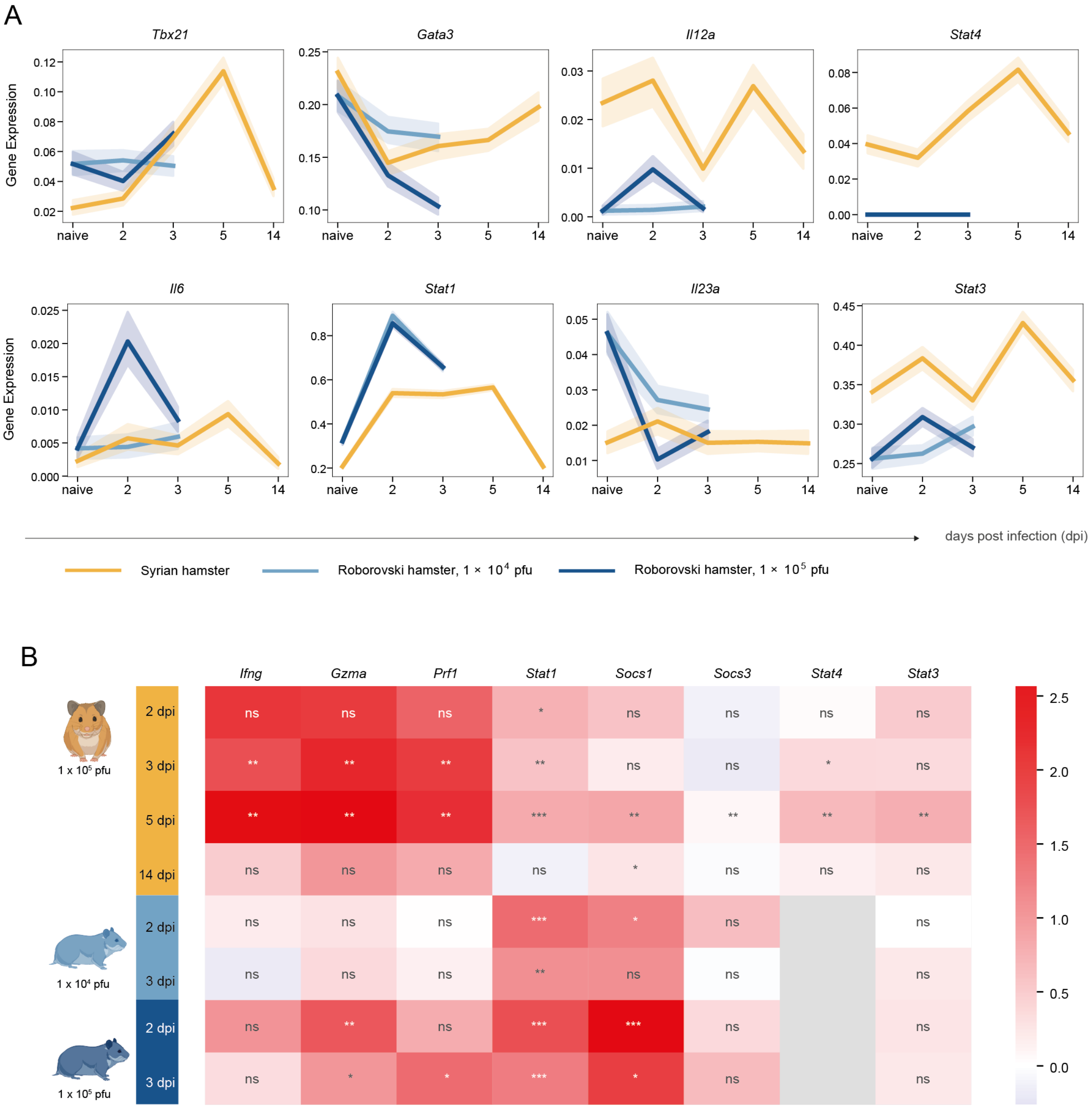


**fig. S3. Selected gene expression along time after infection. (A)** Expression time courses along time after infection per species and virus dose combination for selected immunity marker genes. **(B)** Sample-wise L2FCs of normalized expression with respect to corresponding uninfected samples for selected genes after pseudobulking T cell, NK cell and ILC2 clusters. P-values of t-test for difference in significance across samples (n=3) marked with * < 0.05, ** < 0.01 or ns: not significant.

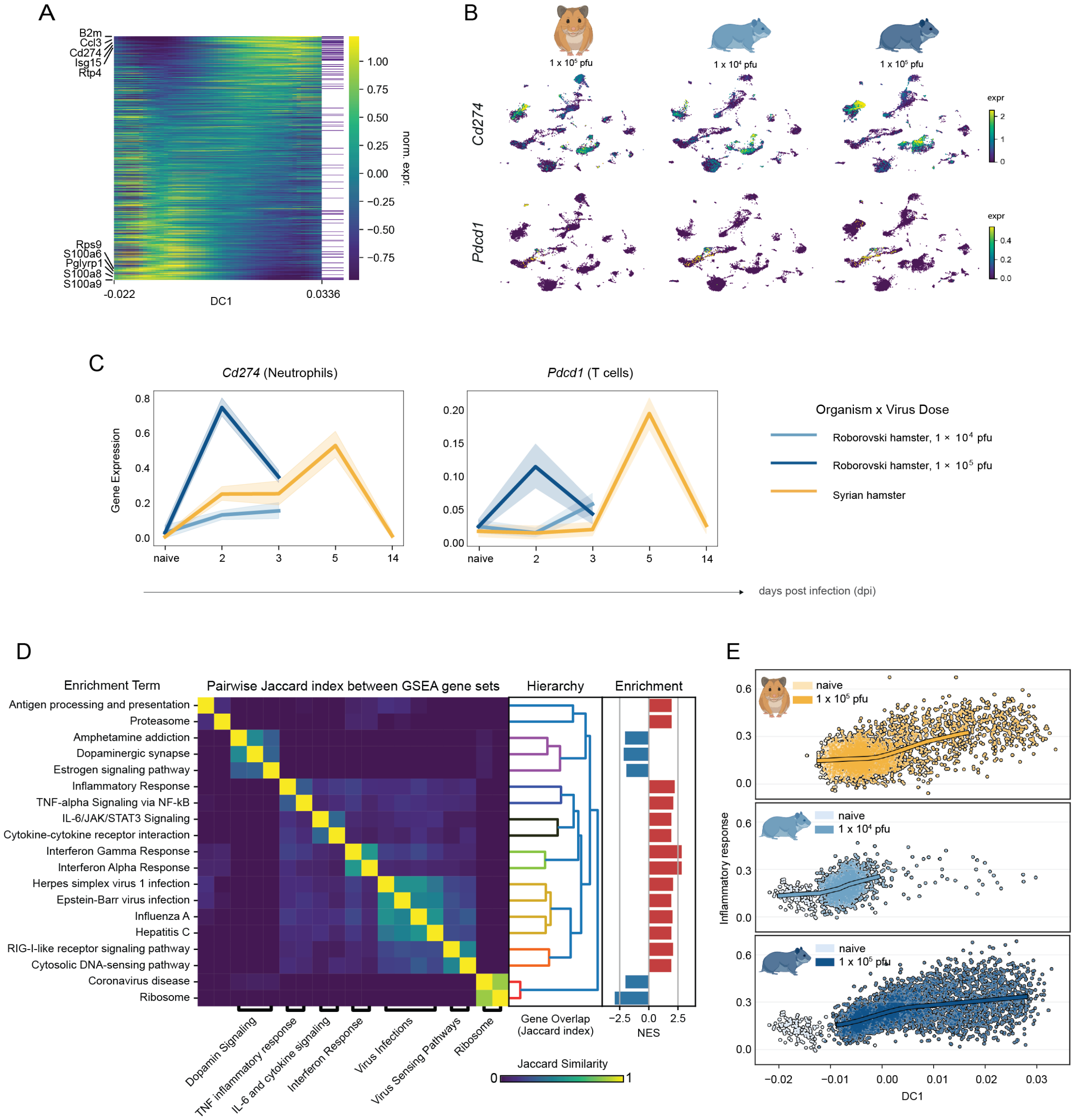


**fig. S4. Supporting analyses for neutrophil response to infection. (A)** Expression of top 100 (anti-) correlating genes along DC1 with genes belonging to the inflammatory response gene set indicated by a purple line in the right column of the heatmap. **(B)** UMAP embeddings of all cells colored by expression of *CD274* (top) and *Pdcd1* (bottom), split by hamster type and virus dose. **(C)** Gene expression trends of *CD274* in neutrophils and *Pdcd1* in T cells along time before and after infection per hamster type and virus dose. Areas around curves are 95% confidence intervals by bootstrapping. **(D)** Pairwise Jaccard index between leading edge gene sets reported by GSEA on DC1 correlating genes. Jaccard index measures the overlap between sets; a value of 1 corresponds to perfect overlap. Resulting hierarchy and associated normalized enrichment scores from GSEA adjacent to the right. Bottom: annotation of heatmap aggregates similar gene sets with high overlap to reduce redundancy. **(E)** Neutrophil DC1 against inflammatory response score split by species and virus dose, colored by infection status. LOESS regression lines shown on top.

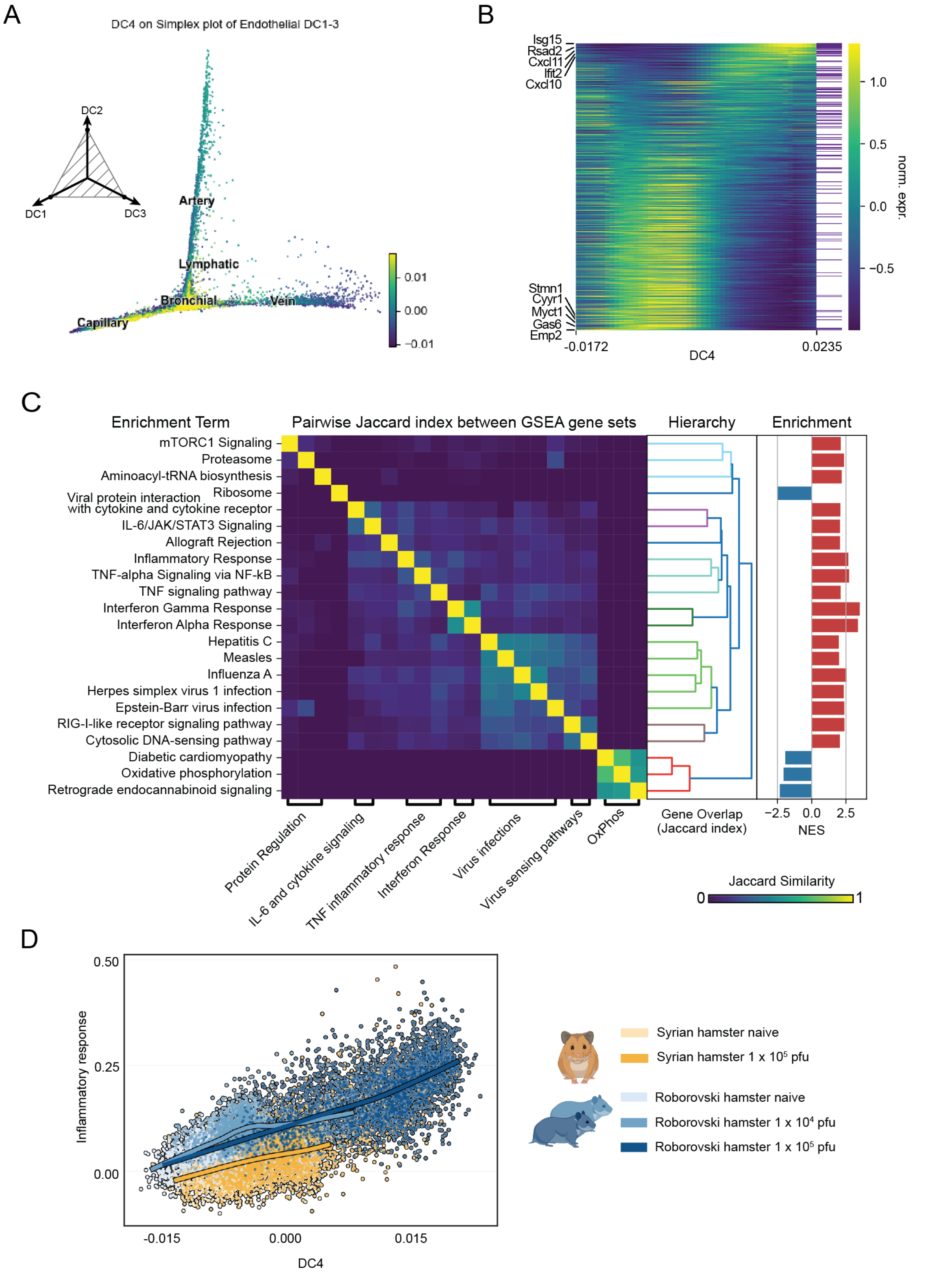
A
DC4 on Simplex plot of Endothelial DC1-3
DC2
DC1
DC3
Artery
Lymphatic
Bronchial
Vein
Capillary
0.01
0.00
−0.01
B
Isg15
Rsad2
Cxcl11
Ifit2
Cxcl10
Stmn1
Cyyr1
Myct1
Gas6
Emp2
-0.0172
DC4
0.0235
1.0
0.5
0.0
−0.5
norm. expr.
C
Enrichment Term
Pairwise Jaccard index between GSEA gene sets
Hierarchy
Enrichment
mTORC1 Signaling
Proteasome
Aminoacyl-tRNA biosynthesis
Ribosome
Viral protein interaction with cytokine and cytokine receptor
IL-6/JAK/STAT3 Signaling
Allograft Rejection
Inflammatory Response
TNF-alpha Signaling via NF-kB
TNF signaling pathway
Interferon Gamma Response
Interferon Alpha Response
Hepatitis C
Measles
Influenza A
Herpes simplex virus 1 infection
Epstein-Barr virus infection
RIG-I-like receptor signaling pathway
Cytosolic DNA-sensing pathway
Diabetic cardiomyopathy
Oxidative phosphorylation
Retrograde endocannabinoid signaling
Protein Regulation
IL-6 and cytokine signaling
TNF inflammatory response
Interferon Response
Virus infections
Virus sensing pathways
OxPhos
Gene Overlap (Jaccard index)
−2.5 0.0 2.5
NES
Jaccard Similarity
0
1
D
0.50
0.25
0.00
Inflammatory response
-0.015
0.000
0.015
DC4
Syrian hamster naive
Syrian hamster 1 x 10^5 pfu
Roborovski hamster naive
Roborovski hamster 1 x 10^4 pfu
Roborovski hamster 1 x 10^5 pfu

**fig. S5. Supporting analyses for endothelial cell response to infection. (A)** Simplex plot from Fig. 5A colored by DC4 value of endothelial cells shows response primarily by capillary and bronchial subtypes. **(B)** Expression of top 100 (anti-) correlating genes along DC1 with genes belonging to the inflammatory response gene set indicated by a purple line in the right column of the heatmap. **(C)** Pairwise Jaccard index between leading edge gene sets reported by GSEA on DC4 correlating genes. Jaccard index measures the overlap between sets; a value of 1 corresponds to perfect overlap. Resulting hierarchy and associated normalized enrichment scores from GSEA adjacent to the right. Bottom: annotation of heatmap aggregates similar gene sets with high overlap to reduce redundancy. **(D)** Endothelial DC4 against inflammatory response score colored by species and virus dose. LOESS regression lines shown on top.
